# Human URAT1/*SLC22A12* gene promoter is regulated by 27-hydroxycholesterol through estrogen response elements

**DOI:** 10.1101/827709

**Authors:** Masaya Matsubayashi, Yoshihiko M. Sakaguchi, Yoshiki Sahara, Hitoki Nanaura, Sotaro Kikuchi, Arvand Ashari, Linh Bui, Shinko Kobashigawa, Mari Nakanishi, Riko Nagata, Takeshi K. Matsui, Genro Kashino, Masatoshi Hasegawa, Shin Takasawa, Masahiro Eriguchi, Kazuhiko Tsuruya, Shushi Nagamori, Kazuma Sugie, Takahiko Nakagawa, Minoru Takasato, Michihisa Umetani, Eiichiro Mori

## Abstract

Elevated levels of uric acid, a metabolite of purine in humans, is related to various diseases, such as gout, atherosclerosis and renal dysfunction. The excretion and reabsorption of uric acid to/from urine is tightly regulated by uric acid transporters. The amino acid sequences of uric acid reabsorption transporters, URAT1/*SLC22A12*, OAT4/*SLC22A11*, and OAT10/*SLC22A13*, share closer phylogenic relationship, whereas the gene promoter sequences are distant phylogenic relationship. Through the single-cell RNA-sequencing analysis of an adult human kidney, we found that only a small number of cells express these transporters, despite their role in the regulation of serum uric acid levels. Transcriptional motif analysis on these transporter genes, revealed that the URAT1/*SLC22A12* gene promoter displayed the most conserved estrogen response elements (EREs) among the three transporters. The endogenous selective estrogen receptor modulator (SERM) 27-hydroxycholesterol (27HC) had positive effects on the transcriptional activity of URAT1/*SLC22A12*. We also found that 27HC increased the protein and gene expression of URAT1/*SLC22A12* in mouse kidneys and human kidney organoids, respectively. These results strongly suggest the role of 27HC for URAT1/*SLC22A12* expression in renal proximal tubules and upregulation of serum uric acid levels and also show the relationship between cholesterol metabolism and serum uric acid regulation.

**Significance Statement:** The elevated levels of serum uric acid cause various diseases, and the excretion/reabsorption of uric acid to/from urine is tightly regulated by the uric acid transporters. We found that despite the role in serum uric acid regulation, only a small number of cells express URAT1/*SLC22A12*. We also found that URAT1/*SLC22A12* gene promoter region has effective estrogen response elements, and endogenous selective estrogen receptor (ER) modulator 27-hydroxycholesterol (27HC) increased URAT1/*SLC22A12* expression in the mice kidneys and human kidney organoids. These suggest that 27HC increases URAT1/*SLC22A12* expression and upregulate serum uric acid levels. Since 27HC connects cholesterol metabolism, our study indicates the important link between cholesterol metabolism and serum uric acid regulation, and also provides a novel therapeutic approach to hyperuricemia.

## Introduction

Elevated serum uric acid levels result in various diseases such as gout, renal disease, and cardiovascular disease (1–5). Serum uric acid levels and prevalence of hyperuricemia increased with aging and also in postmenopausal women (6–8). Increased uric acid also triggers endothelial dysfunction, which leads to metabolic syndrome and cardiovascular diseases (9–14). In the other hand, abdominal obesity, atherogenic dyslipidemia, insulin resistance, aging and hormonal imbalance are the risk factors for metabolic diseases, and patients with metabolic syndrome can have hyperuricemia and hypercholesterolemia (15–18), suggesting the relationship in the regulation of uric acid and cholesterol. However, its precise mechanism has been unknown.

Uric acid is produced in the liver by xanthine oxidoreductase (XOR) and excreted to urine. The excretion of uric acid is regulated by several uric acid transporters, such as urate transporter 1 (URAT1)/*SLC22A12*, ATP-binding cassette sub-family G member 2 (ABCG2)/*ABCG2* and glucose transporter 9 (GLUT9)/*SLC2A9* (19–22). In the proximal tubular (PT) cells, apical urate transporters form a super-molecular structure called urate transportsome, which are organized by a PDZ scaffold (23). Uric acid reabsorption transporters play an important role in regulating serum uric acid levels, and inhibitors of these transporters such as lesinurad and dotinurad attenuate hyperuricemia (24–27).

Previously, we found 27-hydroxycholesterol (27HC) function as an endogenous selective estrogen (ER) modulator (SERM) and promotes atherosclerotic lesion development in an ER-depending manner (28). While estrogen acts as an agonistic and antagonistic on ER in a tissue- and cell-context manner (29, 30). Serum 27HC and cholesterol levels are positively correlated, and serum 27HC levels are also elevated with particularly after 30 years old (29, 31–33),. Furthermore, circulating 27HC levels in premenopausal women are lower than in men, and are increased after menopause (29, 31). In this study, we examined the role of 27HC in renal function, and found that 27HC activates URAT1/SLC22A12 expression via ER using single-cell RNA-sequencing (scRNA-seq), gene promoter analysis, cell culture assays, histology, and kidney organoid assays.

## Results

### Uric acid reabsorption transporters are expressed in kidney PT cells

To analyze the transcription of uric acid transporters at single-cell resolution, we performed the analysis on publicly available scRNA-seq data of human adult kidneys (34) (Fig. 1 and S1). We found that uric acid reabsorption transporters OAT4/*SLC22A11*, URAT1/*SLC22A12*, and OAT10/*SLC22A13*, displayed similar transcription patterns to the markers of proximal tubular cells, cubilin (CUBN), low density lipoprotein-related protein 2 (LRP2), and solute carrier family 5 member 12 (SLC5A12). (Fig. 1C).

**Figure 1.**
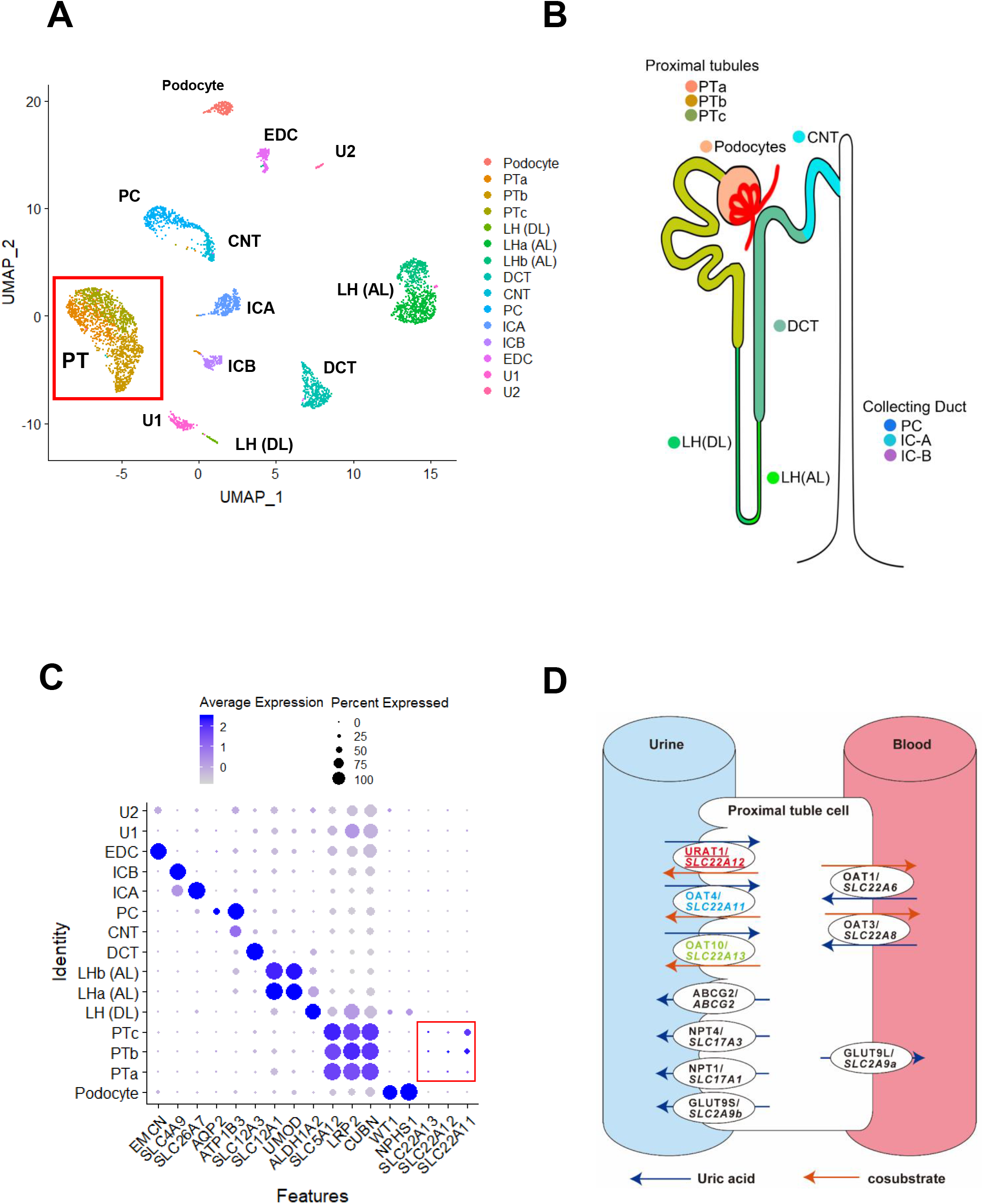
scRNA-seq analysis of human adult kidney. (A) Unsupervised clustering of single-cell RNA-sequence (scRNA-seq) of adult human kidney identified 15 distinct cell types, including podocytes, tubular cells and endothelial cells. (B) Schematic diagram of nephron and connecting duct. Each part shares the same color code as in (A). (C) The dot-plot of nephron individual markers and uric acid reabsorption transporters. PT: proximal tubule, LH (DL): loop of Henle (descending loop), LH (AL): loop of Henle (ascending loop), DCT: distal convoluted tubule, CNT: connecting tubule, PC: principal cell, ICA: intercalated cell type-A, ICB: intercalated cell type-B, EDC: endothelial cell, U: undefined. (D) Schema of the uric acid transporters in the proximal tubular cells. The three reabsorption transporters are displayed in color.

### URAT1, OAT4, and OAT10 are expressed only limited cells in the PT

After glomerular filtration, 90% uric acid is reabsorbed (35). Considering the importance of uric acid efflux/reabsorption in the PT region, we performed scRNA-seq analysis on the distribution of uric acid transporters. Although the expression of secretion uric acid transporters was abundant except for ABCG2/ABCG2 in the PT region (Fig. S4), only a small number of cells express reabsorption transporters URAT1/*SLC22A12*-, OAT4/*SLC22A11-*, or OAT10/*SLC22A13*-positive cells were present in only 9.2%, 26.5%, or 2.1%, respectively, in the PT cells (Fig. 2A-F). Our observations suggest a specific transcriptional regulation that is involved in the expression of uric acid reabsorption transporters.

**Figure 2.**
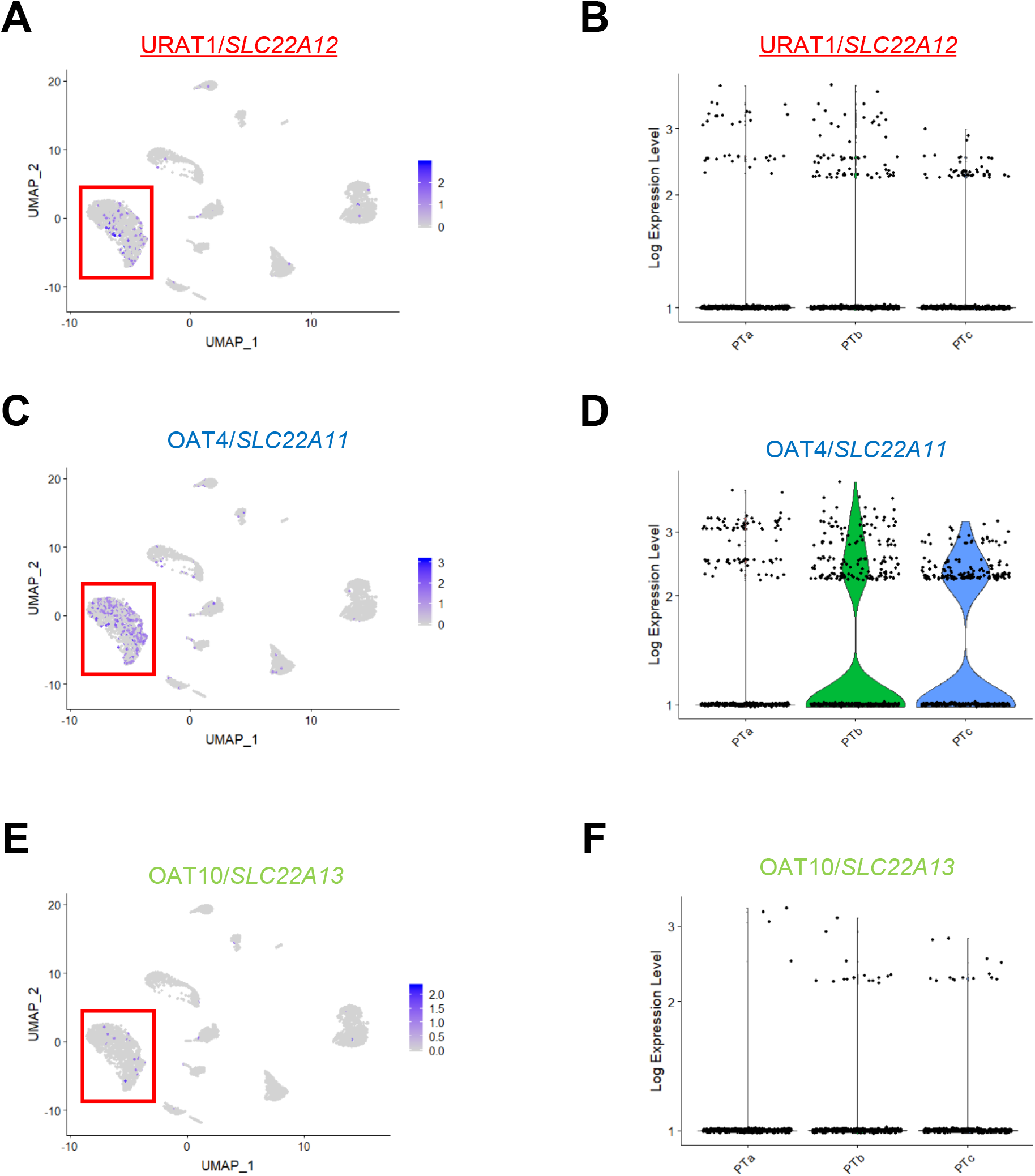
Gene expression of the three uric acid reabsorption transporters at single cell resolution. (A-F) The gene expression level and distribution of URAT1/*SLC22A12* (A, B), OAT4/*SLC22A11* (C, D) and OAT10/*SLC22A13* (E, F). Left panels (A, C, E): feature-plot. Right panels (B, D, F); violin-plot.

### URAT1 is uniquely regulated from other reabsorption uric acid receptors

To investigate the factors that regulate the three reabsorption transporters, URAT1/*SLC22A12*, OAT4/*SLC22A11*, and OAT10/*SLC22A13*, we first analyzed the correlation of their gene expression. Among these genes, OAT4/*SLC22A11* and OAT10/*SLC22A13* showed relatively correlated gene expression patterns (Fig. 3A-C). However, URAT1/*SLC22A12* gene expression showed less correlation with the other reabsorption transporters, suggesting that URAT1/*SLC22A12* expression may have different regulation from the regulation of the other two transporters. Next, we compared the gene promoter region sequences. The promoter sequence of URAT1/*SLC22A12* was distant phylogenic relationship with OAT4/*SLC22A11* or OAT10/*SLC22A13*, further supporting the unique transcriptional regulation of this gene from the other two (Fig.3D). We also analyzed the amino acid sequences of the three reabsorption transporters share closer phylogenic relationship (Fig.3E and S2A). In contrast, from structural prediction, phenylalanine 365 (F365) in URAT1/*SLC22A12* has been previously found to be involved in the recognition of uric acid (36), while the conserved site in the other *SLC22A* transporters is tyrosine, suggesting that URAT1/*SLC22A12* has a higher binding affinity for uric acid (36) (Fig. S2B). These results suggest that the gene expression of URAT1/*SLC22A12* is uniquely regulated from other reabsorption uric acid transporters.

**Figure 3.**
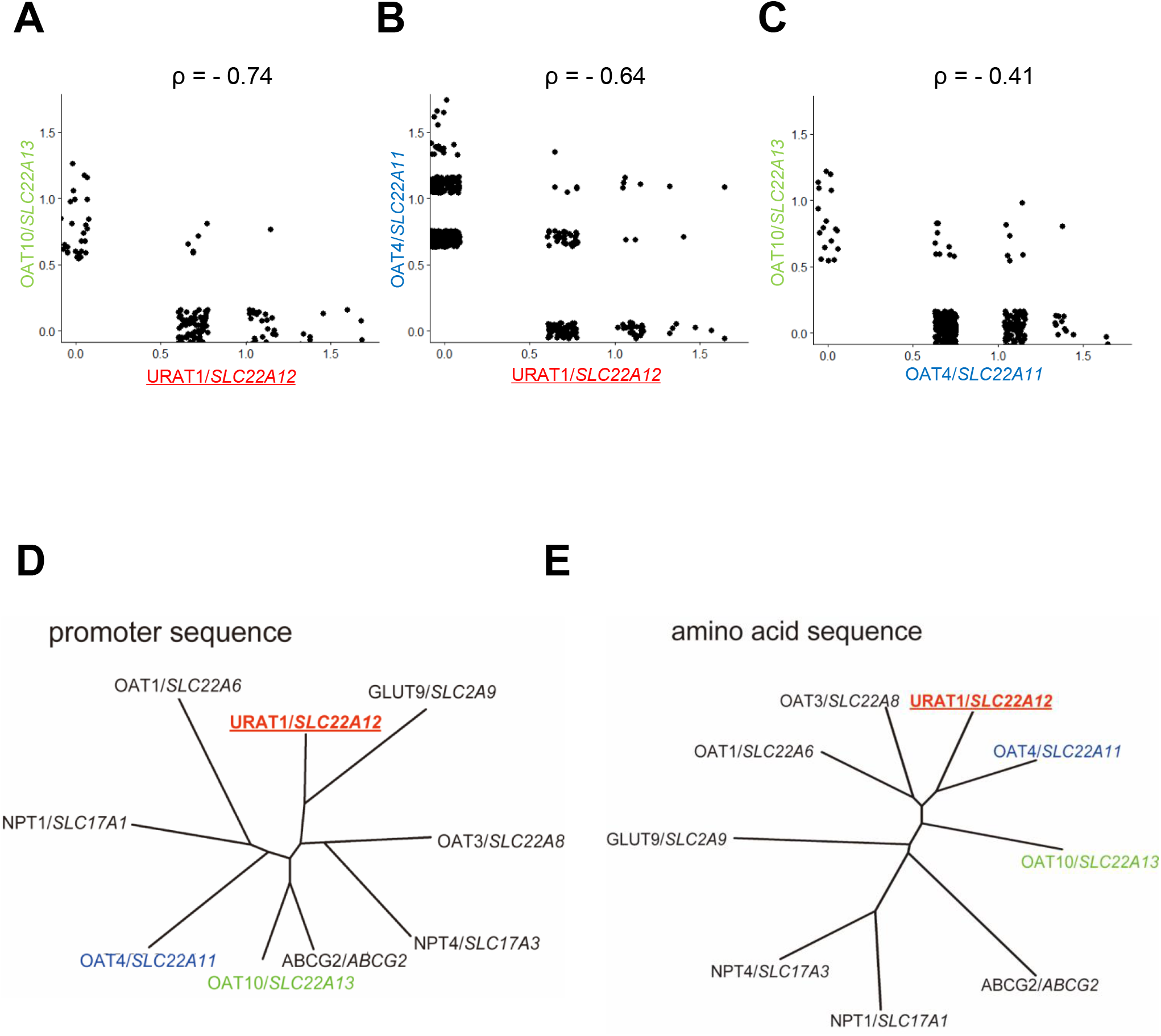
Co-expression analysis of the three reabsorption transporters and the difference and similarity in the amino acid sequences and gene promoter sequences of their transporters. (A-C) Scatter plots show the co-expression of the three uric acid reabsorption transporters. Phylogenic trees of the uric acid transporters (D) promoter region sequences and (E) amino acid sequences. The three reabsorption transporters are displayed in color.

### URAT1/SLC22A12 is regulated by 27HC through ERE gene promoter sequences

To find the key factors for the different transcriptional regulation of reabsorption uric acid transporters, we further analyzed their gene promoter regions. Since increased uric acid is associated with metabolic condition and sex hormone status, we focused whether the gene promoter region has transcription factor motifs related to metabolic or sex hormone-regulated transcription factors. We found that the URAT1/SLC22A12 gene promoter region had nine ERE (estrogen response element) sequences and also the highest ERE value among these three transporter genes (Fig. 4A, 4B and S3). ER binds to EREs and regulate the transcription. ESR1, which encodes estrogen receptor α, but not ESR2, was abundant in the PT cells (Fig. S5A) (34). Thus, we investigated the regulation of URAT1/*SLC22A12* through the identified EREs. To investigate which ERE regulates the URAT1/*SLC22A12* transcription activity responding to ERs, we designed a construct that indicates URAT1/*SLC22A12* promoter activity by luciferase expression (Fig. 4C and D).

**Figure 4.**
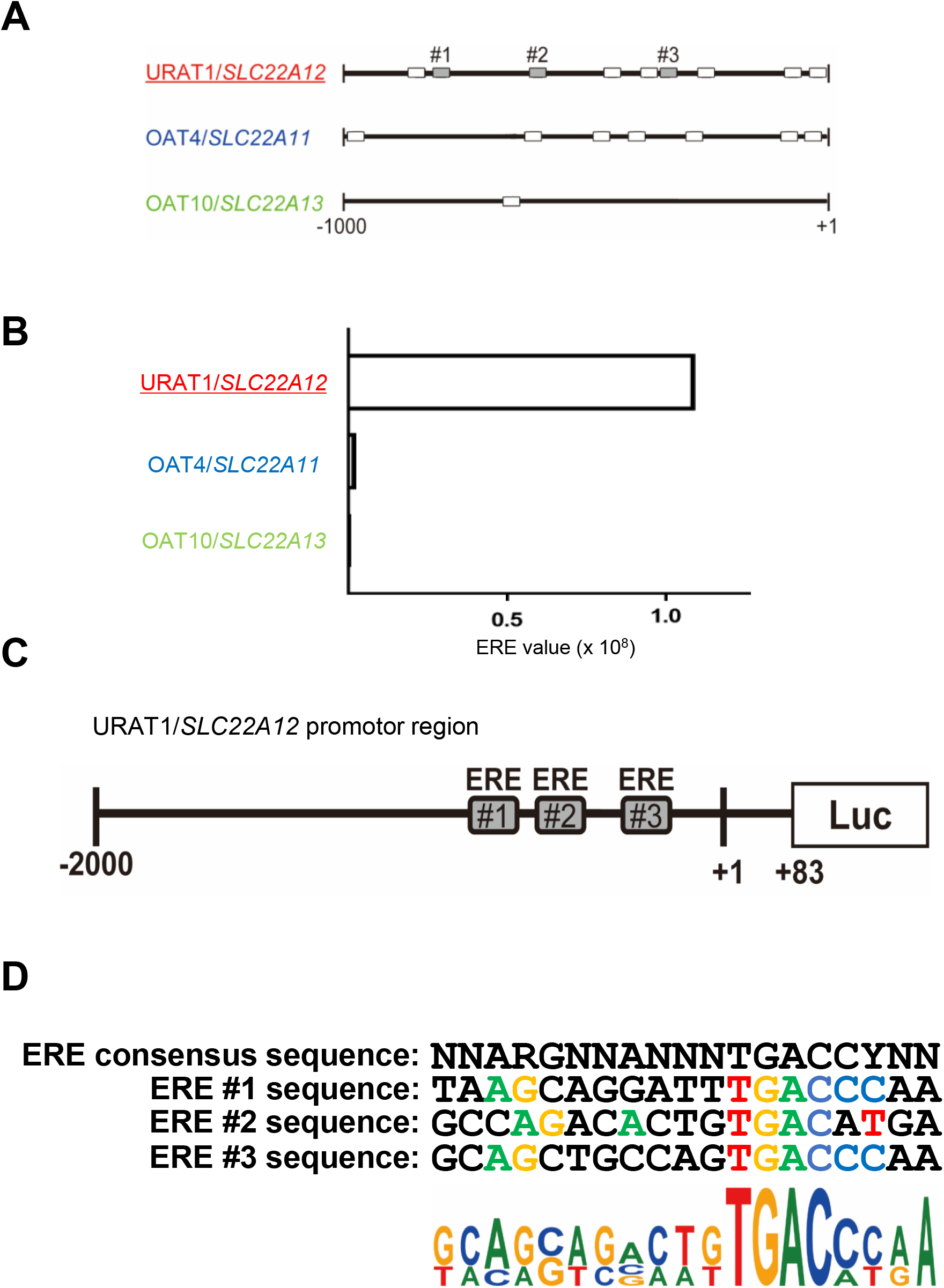
ERE score of the three reabsorption transporters and schematic diagram of URAT1/*SLC22A12* promoter region. (A) A diagram of the promoter region: white boxes show ERE sequence, and three grey boxes of URAT1/*SLC22A12*, #1-#3, show ERE sequence having the ERE score > 0.8. (B) A bar plot showed ERE values of the three uric acid reabsorption transporters promotor region. (C) Schematic representation of human URAT1/*SLC22A12* promoter luciferase reporter constructs. ERE#1, #2 and #3 displays the same site of Fig4A. (D) Consensus sequence of ERE is presented at the top. The others are signal sequences of the predicted ERE. A sequence logo of the three EREs.

Since there is a correlation between serum uric acid levels and metabolic condition, and also 27HC act as an endogenous SERM and regulates transcriptional activities through EREs (33, 37), we examined the effects of 27HC on URAT1/*SLC22A12* gene promoter activity. Gene promoter activity of URAT1/*SLC22A12* was increased by 27HC in HepG2 cells (Fig. 5A). In contrast, 3β-hydroxy-5-cholestenoic acid, a metabolite of 27HC, did not increase URAT1/SLC22A12 gene promoter activity (Fig. 5B), suggesting that 27HC itself, upregulates URAT1/*SLC22A12* gene transcription. To further determine the impacts of 27HC in URAT1/*SLC22A12* promoter region, we treated cells with an ER antagonist, ICI 182,780 (38). ICI 182,780 alone decreased the URAT1/*SLC22A12* gene promoter activity, indicating that it inhibited gene promoter upregulation through endogenous factors (Fig. 5C). The effect of 27HC was suppressed by a co-treatment with ICI 182,780 (Fig. 5D), suggesting that 27HC increased URAT1/*SLC22A12* gene transcription via ER activity. Importantly, ERE mutation decreased the baseline of gene promoter activity (Fig. 5E), indicating that ERE sequence plays a major role to activate URAT1/*SLC22A12* transcription. ERE mutation also inhibited gene promoter upregulation of 27HC (Fig. 5E). These data showed that human URAT1/*SLC22A12* gene promoter is regulated by 27HC via ER through estrogen response elements.

**Figure 5.**
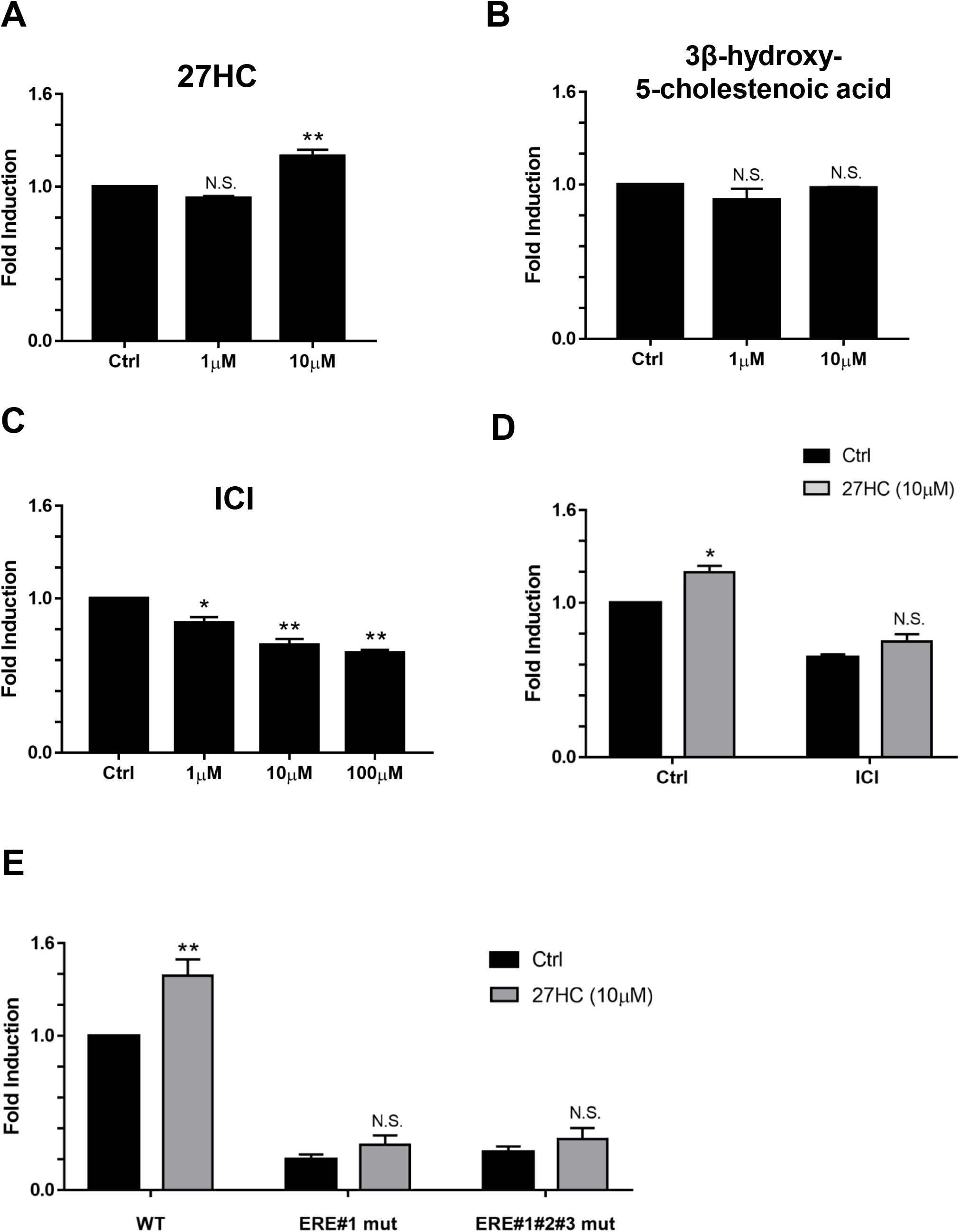
27HC increased URAT1/SLC22A12 gene promoter activity and ER antagonist inhibited the activity through the ERE sequence. (A) The exposure of 27HC for 24 hours showed the effect on URAT1/*SLC22A12* promoter activity in HepG2 cell with stable transcription of the URAT1/*SLC22A12* promoter. The results of one-way ANOVA followed by Dunnett test were as follows: 27HC: *P* = 0.0007, N.S. *P* > 0.05 compared with control, ** *P* < 0.01 compared with control. (B) 3β-hydroxy-5-cholestenoic acid, a metabolite of 27HC, showed the effect on the URAT1/*SLC22A12* promoter activity on HepG2 cells with stable transcription of the URAT1/*SLC22A12* promoter. The results of one-way ANOVA followed by Dunnett test were as follows: 3β-hydroxy-5-cholestenoic acid: *P* = 0.2863, N.S. *P* > 0.05 compared with control. (C) HepG2 cells with stable transcription of the URAT1/*SLC22A12* promoter were exposed to ICI 182,780 (ER antagonist). The results of one-way ANOVA followed by Dunnett test were as follows: ICI 182,780: *P*= 0.0001, * *P* < 0.05 compared with control, ** *P* < 0.01 compared with control. (D) HepG2 cells with stable transcription of the URAT1/SLC22A12 promoter were exposed to 27HC combined with or without 100 μM ICI 182,780 for 24 hours. The results of two-way ANOVA followed by Sidak’s multiple comparisons test were as follows: 27HC treatment: F(1,4) = 16.64, *P* = 0.0151; ICI treatment: F(1,4) = 202, *P* = 0.0001; and 27HC treatment x ICI treatment interaction: F(1,4) = 1.604, *P* = 0.2740. * *P* < 0.05 compared with control. (E) The three HepG2 stable cell lines including ERE mutation were exposed to 10μM 27HC for 24 hours. The results of two-way ANOVA followed by Sidak’s multiple comparisons test were as follows: 27HC treatment: F(1,6) = 24.79, *P* = 0.0025; ERE mutation: F(2,6) = 111.4, *P* < 0.0001; and 27HC treatment x ERE mutation interaction: F(2,6) = 6.895, *P* = 0.0279. ** *P* < 0.01 compared with control. Each experiment was performed in independently three times. All data are mean ± SEM.

### 27HC increases URAT1/SLC22A12 expression in the kidney

Finally, to confirm above URAT1/*SLC22A12* transcriptional regulation in organs, we employed mouse kidneys and human kidney organoids. 27HC is metabolized by CYP7B1, and the loss of *Cyp7b1* in mouse results in elevated serum 27HC levels without affecting cholesterol and bile acid levels (39). Therefore, we performed immunohistochemistry using kidneys from *Cyp7b1^-/-^* and wild-type mouse. URAT1/*SLC22A12* protein expression was observed in PT region and, the expression levels increased in male *Cyp7b1^-/-^* mouse (Fig. 6A and B). Interestingly, female *Cyp7b1^-/-^* mouse did not show URAT1/*SLC22A12* expression levels (Fig. S8A and B), suggesting the involvement of sex hormone coregulation. Furthermore, we performed kidney organoid culture from human iPS cells to examine the effect by 27HC on the URAT1/*SLC22A12* expression in human kidneys, because kidney organoids mimic human embryo kidney. 27HC increased URAT1/*SLC22A12* gene expression in the kidney organoids (Fig. 6C), suggesting that its effects on URAT1/*SLC22A12* expression is also observed in human kidneys.

**Figure 6.**
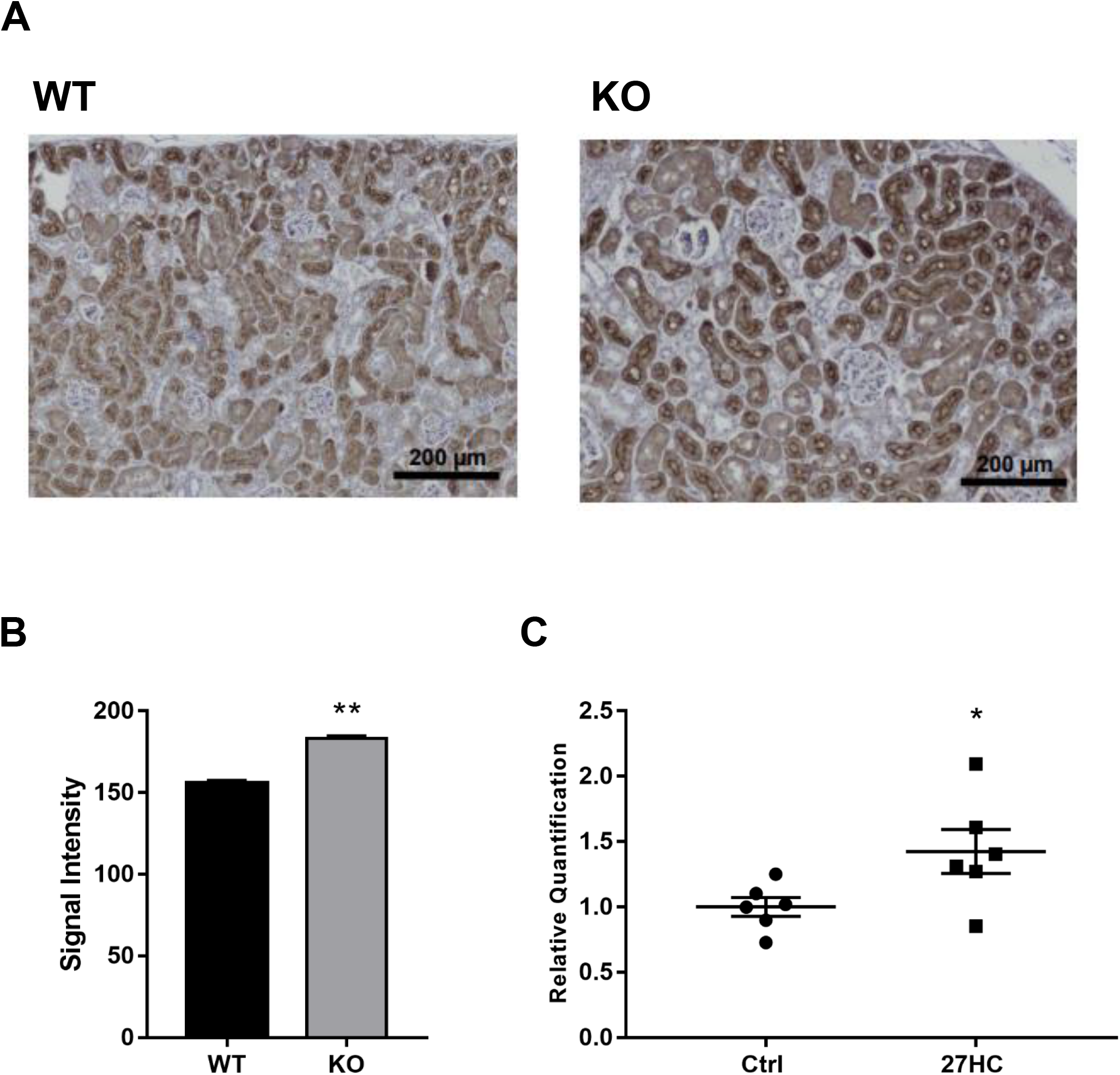
27HC increased URAT1/SLC22A12 expression in the kidneys. (A) Immunohistochemical analysis of the kidney in the male *Cyp7b1^-/-^* mouse. Left panel: wild type mouse, right panel: *Cyp7b1*^-/-^ mouse. Scale bar showed 200μm. (B) We performed image analysis on (A) (n = 240 proximal tubular cells). ** *P* < 0.01 compared with control by student’s t-test. (C) 27HC induced URAT1/*SLC22A12* gene expression in the human kidney organoids. The experiment was performed in six independently experiments. * *P* < 0.05 compared with control by student’s t-test. The data are mean ± SEM.

## Discussion

Our current study demonstrated that 27HC, an endogenous SERM, can increase serum uric acid levels by regulating the expression of URAT1/*SLC22A12* and serum uric acid levels (Fig. S8). A high-fat diets and alcohol consumption are known to increase serum uric acid levels, and these factors are consequently considered to activate xanthine oxidoreductase by increasing purine. Although the expression of URAT1/*SLC22A12* increased in hyperuricemia or obesity in animal models, there was limited information on the mechanism(s) by which URAT1/*SLC22A12* expression is regulated (40–42). URAT1/*SLC22A12* localizes on the apical membrane of proximal tubular cells and is deemed to be involved in both pre-secretary and post-secretary reabsorption (43, 44). The distribution of URAT1/*SLC22A12* on the proximal tubule remains controversial; one report showed its distribution on S1 and S3 segments (43), while another group found the transporter on S1 segments only (44). To transcriptionally characterize URAT1/*SLC22A12*-positive cells, we analyzed human adult kidney scRNA-seq datasets (23) and identified the URAT/*SLC22A12*-positive cells at single cell resolution (Fig. 2A and B). Most of filtered uric acid is reabsorbed from urine, and URAT1/*SLC22A12* contributes to the reabsorption (35). The importance of URAT1/*SLC22A12* was demonstrated by using URAT1/*SLC22A12* inhibitors dotinurad and lesinurad that decreased serum uric acid levels (24–26). Increased expression of URAT1/*SLC22A12* was also been found in pathological rodent model (40–42). Initially, we hypothesized that the risk factor of various diseases would affect URAT1/*SLC22A12* expression and increase the number of URAT1/*SLC22A12*-positive cells. Nonetheless, we found that the number of the URAT1/*SLC22A12*-positive cells was unexpectedly small (9.3%) (Fig. 2A and 2B). These results, together with the abundant expression of uric acid efflux transporters in the PT region, suggest that limited number of cells regulate uric acid reabsorption by changing the expression of URAT1/*SLC22A12*.

URAT1/*SLC22A12* gene promoter has nine EREs. ER regulates the transcriptional of its target gene through EREs. 27HC, an endogenous SERM, promotes atherosclerosis through its activation of proinflammatory processes that are mediated by ER. In this study, we found that 27HC also increases URAT1/*SLC22A12* gene and protein expression. The upregulation URAT1/*SLC22A12* gene promoter activity by 27HC was suppressed by both ICI 182,780, an ER antagonist, and mutation on the ERE sequences on the gene promoter region (Fig. 5D and 5E). The circulating level of 27HC is higher in men than in women and also increased with age (29, 31, 32). Furthermore, in women the serum 27HC levels increase after menopausal that greatly lowers estrogen biosynthesis (31). These trend resembles uric acid levels. In addition, the activity of kidney CYP7B1 decreases with age, which further increases local 27HC levels in the kidney (45). Taken together with our results, it is suggested that the increment of the serum and kidney 27HC levels with age upregulates URAT1/*SLC22A12* transcriptional activity and thereby increases serum uric acid levels (Fig. S9).

The 27HC is the most abundant oxysterol in human circulation, and its circulating levels are elevated with hypercholesterolemia and metabolic or cardiovascular dysfunction such as atherosclerosis. Therefore, the expression of URAT1/*SLC22A12* is elevated in obese and atherosclerosis condition, at least in part, by elevated 27HC levels. URAT1/*SLC22A12* protein expression is increased in mice fed a high-fat diets and also in genetically modified obese (*ob/ob*-) mice (42). Although the previous report did not measure 27HC levels, 27HC may have had an impact on the expression of URAT1/*SLC22A12*. Urinary uric acid excretions were observed lower in obese people compared with people with normal body weight. Additionally, body mass index is positively correlated to serum uric acid levels, suggesting that 27HC and URAT1 protein levels are increased in obese people, leading the increase in serum uric acid levels.

Abnormal levels of uric acid cause various diseases, such as gout, renal disease, and cardiovascular diseases. Based on our findings, it is plausible that elevated 27HC levels induce high serum uric acid levels and promote the related diseases. Our findings in this study highlight the importance of controlling serum uric acid and cholesterol levels, help to understand the correlation between hyperuricemia and metabolic syndrome, and also provide a potential therapeutic approach toward hyperuricemia-related diseases by modifying 27HC levels.

## Materials and Methods

### Single-cell RNA-sequencing analysis

Single-cell transcriptome analysis on the human adult kidney was performed using a dataset of single nucleus RNA-seq from a human adult kidney (GEO: GSE118184) (34). Obtained digital gene expression matrix was processed for quality control, dimensionality reduction and cell clustering using Seurat (https://satijalab.org/seurat/) plug-in of R software (https://www.r-project.org/). Cells were filtered out as low-quality cells judging from unique feature counts and mitochondrial counts. After the filtration, we employed sctransform normalization (46, 47). During the normalization, we removed the confounding sources of variation; mitochondrial mapping percentage, the total number of molecules and cell cycle heterogeneity. Principal component (PC) analysis using the function RunPCA generated clusters was conducted and the first top significant PCs were selected for two-dimensional Uniform Manifold Approximation and Projection (UMAP) with the methods of PCHeatmap, JackStrawPlot and PCElbowPlot. Unsupervised clustering was done by the functions, FindClusters and RunUMAP at a resolution level of 0.8. Each cluster was identified using marker genes reported by a previous report (34). The hierarchical clustering heatmap was generated using heatmap.2 function from the ggplot2 package with the average expression of each cluster against the marker genes. The dot-plot was generated with the average expression of each cluster against the marker genes.

### Alignment of uric acid transporter

We identified the 1000bp upstream sequences in gene promoter region and amino acid sequences of human uric acid transporters (URAT1/*SLC22A12*, OAT4/*SLC22A11*, OAT10/*SLC22A13*, ABCG2/*ABCG2*, NPT1/*SLC17A1*, NPT4/*SLC17A3*, GLUT9/*SLC2A9*, OAT1/*SLC22A6* and OAT3/*SLC22A8*) derived from the NCBI database. Multiple alignment was performed by MUSCLE. Phylogenetic tree was generated by ClustalW 2.0 using the Neighbor-Joining (NJ) method (48, 49) and DRAWTREE in PHYLIP software suite (50).

### Transcription factor (TF) search

We searched TF sites by using TFBIND (http://tfbind.hgc.jp/). Sequence Logo was made by Web Logo 3 (http://weblogo.threeplusone.com/.

### ERE score

ERE score was defined as the calculated consensus score by TFBIND. ERE values were represented as multiplication of the ten-fold ERE scores.

### URAT1/*SLC22A12* promoter cloning

All constructs were designed by SnapGene (GSL Biotech LLC). The human gDNA was isolated from U2OS cells using the QuickExtract™ DNA Extraction Solution (EPICENTRE). From the gDNA, the URAT1/SLC22A12 gene promoter region was amplified using KOD One PCR Master Mix (TOYOBO) or PrimeSTAR^®^ Max DNA Polymerase (Takara) with the following primers; primer F: gctcgctagcctcgaaatcagttccaaagagcctctaaagaag, primer R: ccggattgccaagctagagaggcagctgctcca. After the PCR product was confirmed by the agarose gel electrophoresis, it was purified using QIAquick Gel Extraction Kit (Qiagen). The pGL4.17[luc2/Neo] Vector (Promega) was digested with XhoI and Hind III (TOYOBO). The PCR product was inserted into the linearized pGL4.17[luc2/Neo] vector by In-Fusion reaction using In-Fusion HD Cloning Kit (Takara). The URAT1/*SLC22A12* promoter + pGL4.17 plasmid was transformed into the *E. coli*, Competent Quick DH5α (TOYOBO) on the LB agar plate. A few colonies were picked up and expanded overnight in LB medium with ampicillin (50μg/ml). The plasmid was purified using Wizard® Plus SV Minipreps DNA Purification Systems (Promega). After checking the insert in the plasmid using PCR and electrophoresis, the plasmid sequence was checked by the Macrogen Online Sequencing Order System to confirm the value of the construct.

### ERE mutagenesis

All constructs were designed by SnapGene (GSL Biotech LLC). The ERE sequences of URAT1/SLC22A12 promoter region was amplified to get mutants using PrimeSTAR^®^ Mutagenesis Basal Kit (Takara) with the following primers, ERE#1 primer F: aagcaggattcagtacaagggctcgcagtgcgt, primer R: cgagcccttgtactgaatcctgcttaaaagcca, ERE#2 primer F: ccagacactgcagtatgagaggccatagctgag, primer R: tggcctctcatactgcagtgtctggcctggaac, ERE#3 primer F: cagctgccagcagtacaagcccacacagagact, primer R: tgtgggcttgtactgctggcagctgcccagccc.

### Cell culture and generation of stable cell line

The human hepatocellular carcinoma cell HepG2 was purchased from Cellular Engineering Technologies, Inc. HepG2 was maintained in RPMI1640 medium (Nacalai, Japan). Theses media were supplemented with 10% fetal bovine serum, 100 U/mL penicillin, and 100μg/mL streptomycin. The cells were cultured under a humidified atmosphere containing 5% CO_2_ at 37°C. To generate a stable line, cells were seeded in six-well plate at 1.0 × 10^5^ cells per well and transfected with linearized luciferase expression plasmid fused with URAT1/*SLC22A12* gene promoter region using Lipofectamine 2000 (Invitrogen). At 24 hours after transfection, medium was changed to G418 (800 μg/mL)-containing medium. We selected the stably transfected cells by changing medium every day for 2 weeks.

### URAT1/*SLC22A12* luciferase assay

The luciferase activity was measured in WT and ERE mutant URAT1/*SLC22A12* promoter stably expressing HepG2 cells using One-Glo EX Luciferase Assay System (Promega) according to manufacturer’s instructions. Briefly, 24 hours after seeding stable cells in ninety-six well plates, we exposed cells with 27HC, 3β-hydroxy-5-cholestenoic acid or ICI 182,780 for 24 hours. Then, 80μL of One-Glo EX reagent was added in each well and incubated for three minutes to lyse cells. Finally, luciferase activity was measured with a SpectraMax. All treatment experiments were performed in triplicate.

### Kidney immunohistochemistry

Kidney samples were collected from 12 months old *Cyp7b1^+/+^* and *Cyp7b1^-/-^* mice on the C57BL/6 background. Collected samples were fixed in 4% formalin for 48 hours in 4°C, processed, paraffin embedded, and cut into 6-μm thick sections. After rehydration, the sections were treated in the citrate buffer (pH=6) at 95°C for ten minutes for antigen retrieval, followed by 10% goat serum (Life Technologies) at room temperature for one hour. Then, the sections were blocked using avidin/biotin blocking kit (Vector Laboratories) for 15 minutes, added anti-URAT1 antibody (Milipore Sigma, HPA024575, 1:500), and incubated for overnight. After washing the sections, they were blocked again using 3% hydrogen peroxidase (ThermoFisher Scientific), and incubated with biotinylated goat-anti-rabbit antibody (Vector Laboratories, BA-1000, 1:200) for one hour. The sections were visualized using avidin biotin complex kit (ABC kit, Vector Laboratories) and DAB Quanto chromogen (ThermoFisher Scientific), followed by counterstaining with Harris hematoxylin (ThermoFisher Scientific).

### Image analysis

We performed the image analysis using Image J/Fiji and detected the proximal tubule and measured the max intensity of brush border membrane in the proximal tubular cells. 240 points per image were measured in a blind-manner.

### Kidney organoid culture

Organoids differentiation induced from human pluripotent stem cells. Kidney organoids were generated from human induced pluripotent stem cell line CRL1502 according to our protocol (51) with some modification. Briefly, human iPS cells were treated with 8 μM CHIR99021 in APEL2 (STEMCELL Technologies) supplemented with 1% Protein Free Hybridoma Medium II (PFHM II, GIBCO) (basal medium) for 5 days, followed by FGF9 (200 ng/ml) and heparin (1 μg/ml) for another 2 days. Then, cells were collected and dissociated into single cells using trypsin. 0.25 x 10^6^ of collected cells were spun down at 400 x g for 2 min to form a pellet and pellets were transferred onto a Transwell of 0.4 μm pore polyester membrane (#3450 Corning) to culture in liquid-air interfaces. Pellets were treated with 10 μM CHIR99021 in basal medium for 1hour, and then cultured with FGF9 (200 ng/ml) and heparin (1 μg/ml) for another 5 days, followed by another 13-15 days in basal medium. Organoids were treated with 10 μM 27HC for last 2 days before sampling (day25 – 27 of differentiation).

### Quantitative RT-PCR

Total RNA was extracted from organoids using Nucleo Spin (MACHEREY NAGEL) and cDNA was synthesized from 500 ng of total RNA using PrimeScript™ RT Master Mix (TAKARA BIO). We performed quantitative reverse-transcription PCR (qRT-PCR) with SsoAdvanced Universal SYBR Green Supermix (Bio-Rad Laboratories, Hercules, USA) and specific primers for qRT-PCR. We also run the reaction in triplicate and normalized the transcription of each gene to the mean of β-actin as a housekeeping gene and analyzed using the ΔΔCT method by StepOne Software Ver2.3. The primer sequences for qRT-PCR were designed by Bio-Rad (URAT1/*SLC22A12*; qHsaCID0011483, β-actin; qHsaCED0036269).

### Statistics and data analysis

The result of the luciferase assay and quantitative RT-PCR was represented by the ratio of control mean. The values obtained are described as means ± SEM. We performed student’s t-test, a one-way ANOVA followed by Dunnett test or two-way ANOVA followed by Sidak’s multiple comparisons test to determine the significance among groups. All statistical analysis was performed by GraphPad Prism7. scRNA-seq measures transcripts from both cytoplasm and nucleus, whereas single nucleus RNA-seq measures only nuclear transcripts. Nuclei contains only a fraction of total cell RNA. Although nuclear and cytoplasmic mRNAs are highly correlated (52), some protein-coding mRNAs are retained in the nucleus (53). Despite these differences, single-cell and snRNA-seq datasets predict cell types comparably with high concordance (54, 55).

## Acknowledgements

The authors thank Keren-Happuch E Fan Fen for her critical reading of the manuscript and Aya Shimada for assisting our study. This work was supported by grants from JSPS KAKENHI [JP17H07031 to E.M., JP16K19836 to S.K., JP18H06202 to H.N., JP19K16925 to T.K.M], Takeda Science Foundation to E.M. and T.K.M., Kanzawa Medical Research Foundation to E.M., Uehara Memorial Foundation to E.M., Nakatomi Foundation to E.M., Konica Minolta Science and Technology Foundation to E.M., Naito Foundation to E.M., MSD Life Science Foundation to E.M., Mochida Memorial Foundation for Medical and Pharmaceutical Research to E.M., SENSHIN Medical Research Foundation to E.M., Terumo Foundation for Life Sciences and Arts to E.M., Nara Kidney Disease Research Foundation to E.M., Novartis Research Grants to E.M., K.S. and N.E., Nara Medical University Grant-in-Aid for Collaborative Research Projects to K.S., Nara Medical University Grant-in-Aid for Young Scientists to T.K.M., Sumitomo Dainippon Pharma Research Grant to T.K.M., National Institutes of Health grant HL127037 to M.U., and by unrestricted funds provided to E.M. from Dr. Taichi Noda (KTX Corp., Aichi, Japan) and Dr. Yasuhiro Horii (Koseikai, Nara, Japan).

## Supplemental Figure Legend

**Figure S1. scRNA-seq analysis of human adult kidney.** (A) Clusters were visualized by UMAP. (B) Heatmap of transcription of top-5 marker genes defining clusters shown in Fig. 1A.

**Figure S2. Amino acid sequence alignment of human uric acid transporters in the proximal tubular cells.** (A) The alignment was made using MUSCLE. The red shades indicate conserved amino acids. (B) The phenylalanine at the 365 and 366 site of URAT1 could confer the specificity for uric acid of URAT1/*SLC22A12*, and differs from other OATs which with tyrosine at the same site.

**Figure S3. Promoter sequence alignment of human uric acid reabsorption transporters.** (A) The alignment was made using MUSCLE. (B) ERE position and score in uric acid reabsorption transporters promoter.

**Figure S4. The gene expression of uric acid transporter at single-cell resolution.** (A) NPT1/SLC17A1, (B) NPT4/SLC17A3, and (C) ABCG2/ABCG2 efflux uric acid into urine from the proximal tubular cells. (D) GLUT9/*SLC2A9* transports from the cells into urine or blood. (E) OAT1/SLC22A6 and (F) OAT3/SLC22A8 import uric acid from blood into the cells. Left panel: feature-plot. Right panel; violin-plot.

**Figure S5. Expression of the genes related to estrogen metabolism and uric acid at single-cell resolution.** (A) *ESR1* and (B) *ESR2* encode estrogen receptor α and β. (C) *HSD17B11* is responsible for the interconversion of estrone (E1) and estradiol (E2). (D) *SULT1E1* transforms estrone sulfate into estrone (E1). (E) STS converts estrone (E1) into estrone sulfate. (A)-(E) Left panel: feature-plot. Right panel; violin-plot.

**Figure S6. Expression of the genes related to 27HC metabolism at single-cell resolution.** (A) *CYP27A1* synthesizes 27HC from cholesterol. (B) *CYP7B1* metabolizes 27HC. Left panel: feature-plot. Right panel; violin-plot.

**Figure S7. Expression of URAT1/*SLC22A12* in mouse and human kidney models with elevated 27HC.** (A) Immunohistochemical analysis of the kidney in the female *Cyp7b1^-/-^* mouse. Left panel: wild type mouse, right panel: *Cyp7b1^-/-^* mouse. Scale bar showed 200μm. (B) We performed the image analysis on (A) (n = 240 proximal tubular cells). (C) Procedures of human kidney organoids with 27HC. All data are mean ± SEM.

**Figure S8. Graphical abstract.** 27HC binds to ER and activate URAT1/*SLC22A12* transcription. This results in the increase of URAT1/*SLC22A12* in the apical membrane, and uric acid is transported into the cells, followed by increased serum uric acid levels. The red arrow demonstrates an upregulation of URAT1/*SLC22A12* transcription. Blue arrows show the movement of uric acid.

