## Supplement figure for "Human URAT1/*SLC22A12* gene promoter is regulated by 27-hydroxycholesterol through estrogen response elements"

**A**

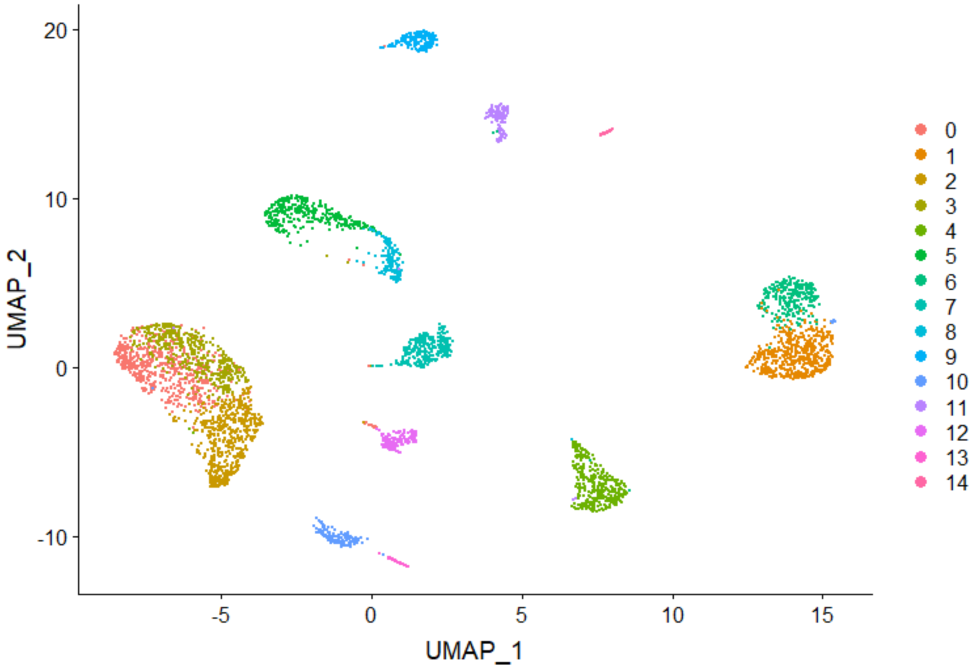
20
10
0
-10
UMAP_2
-5
0
5
10
15
UMAP_1

**B**

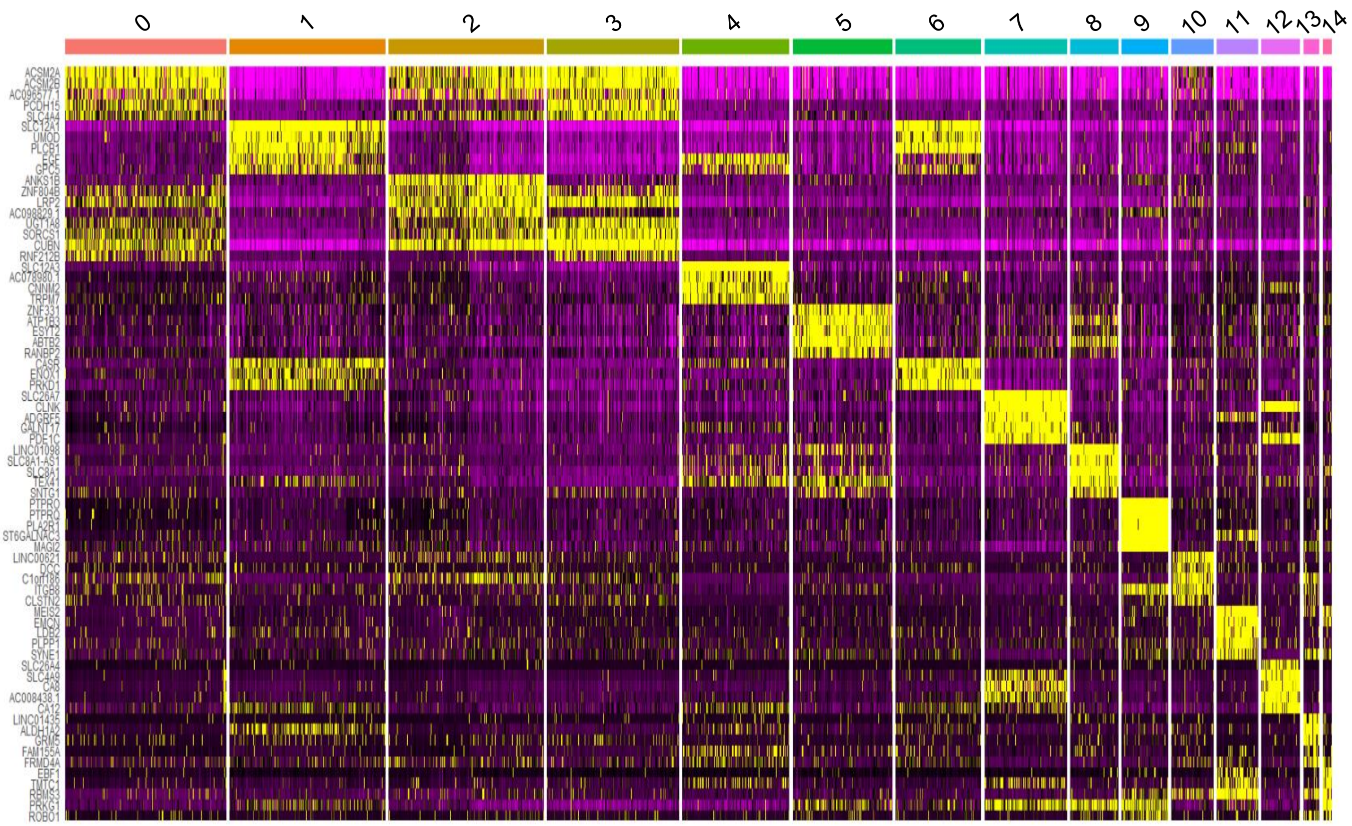
ACSM2A
ACSM2B
AC096577.1
PCDH15
SLC4A4
SLC12A1
UMOD
PLCB1
EGF
GPC5
ANKS1B
ZNF804B
LRP2
AC098829.1
UGT1A8
SORCS1
CUBN
RNF212B
SLC12A3
AC078980.1
CNNM2
TRPM7
ZNF331
ATP1B3
ESYT2
ABTB2
RANBP2
CASR
ENOX1
PRKD1
SLC26A7
CLNK
ADGRF5
GALNT17
PDE1C
LINC01098
SLC8A1-AS1
SLC8A1
TEX41
SNTG1
PTPRO
PTPRQ
PLA2R1
ST6GALNAC3
MAGI2
LINC00621
DCC
C1orf186
ITGB8
CLSTN2
MEIS2
EMCN
LDB2
PLPP1
SYNE1
SLC26A4
SLC4A9
CA8
AC008438.1
CA12
LINC01435
ALDH1A2
GRM5
FAM155A
FRMD4A
EBF1
TMTC1
RBMS3
PRKG1
ROBO1

Matsubayashi, Sakaguchi et al. Supplemental Figure 2

**A**

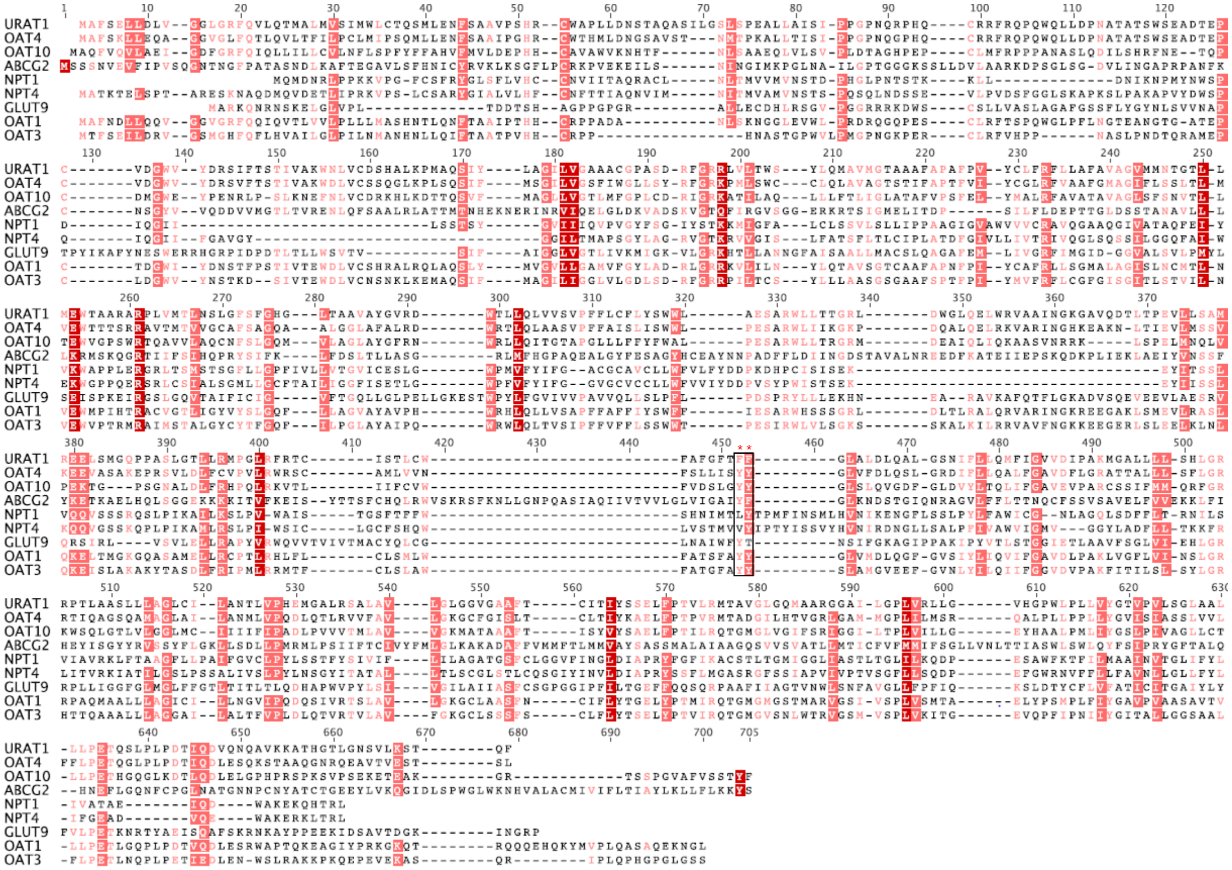

**B**

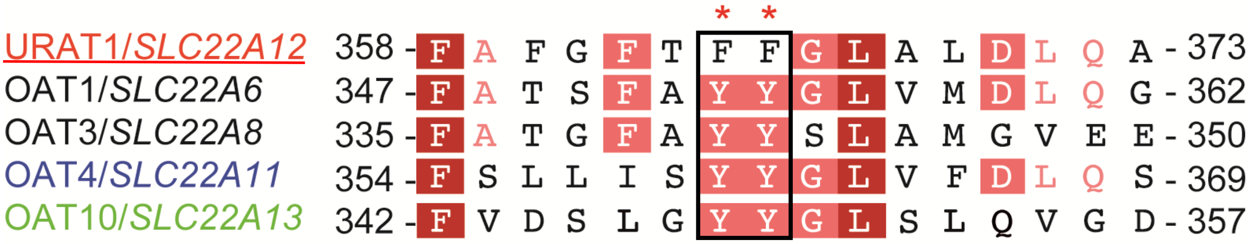

### Matsubayashi, Sakaguchi et al. Supplemental Figure 3

## A

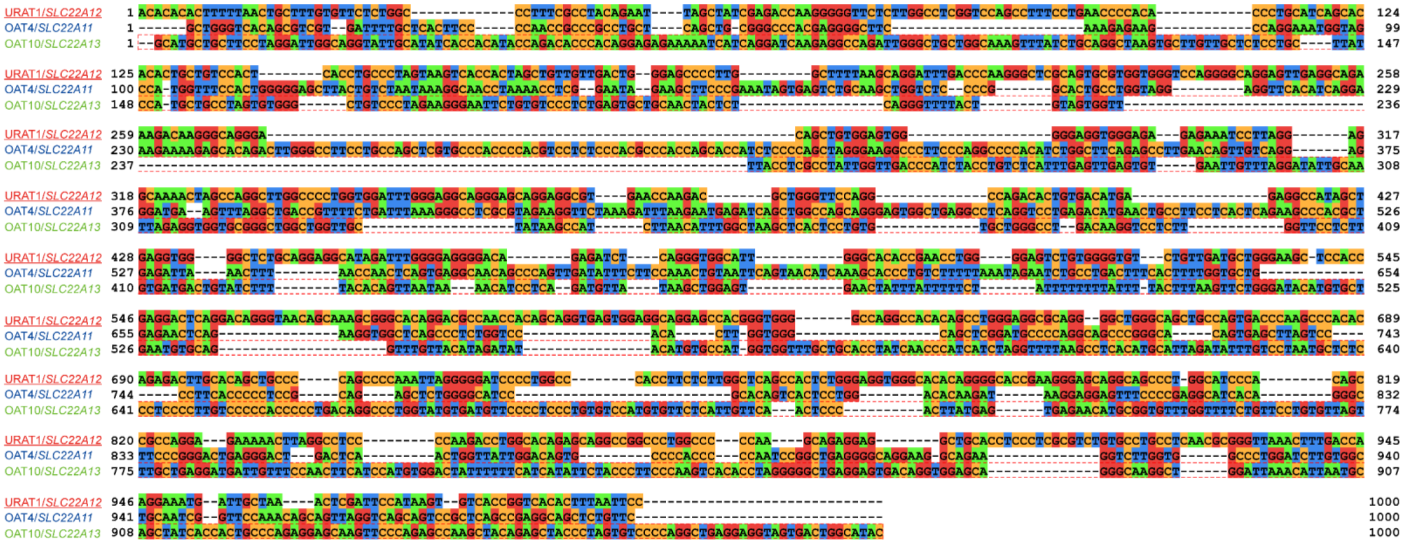

## B

URAT1/*SLC22A12*

| | start | stop | score |
|---|---|---|---|
| ERE1 | -849 | -831 | 0.748099 |
| ERE2 | -805 | -787 | 0.827709 |
| ERE3 | -603 | -585 | 0.859078 |
| ERE4 | -448 | -430 | 0.748337 |
| ERE5 | -371 | -353 | 0.743346 |
| ERE6 | -337 | -319 | 0.854563 |
| ERE7 | -252 | -234 | 0.735029 |
| ERE8 | -71 | -53 | 0.748099 |
| ERE9 | -19 | -1 | 0.780181 |

OAT4/*SLC22A11*

| | start | stop | score |
|---|---|---|---|
| ERE1 | -998 | -980 | 0.865019 |
| ERE2 | -620 | -602 | 0.789686 |
| ERE3 | -472 | -454 | 0.734791 |
| ERE4 | -405 | -387 | 0.737167 |
| ERE5 | -279 | -261 | 0.855513 |
| ERE6 | -89 | -71 | 0.747861 |
| ERE7 | -37 | -19 | 0.730989 |

OAT10/*SLC22A13*

| | start | stop | score |
|---|---|---|---|
| ERE1 | -662 | -644 | 0.743108 |

Matsubayashi, Sakaguchi et al. Supplemental Figure 4

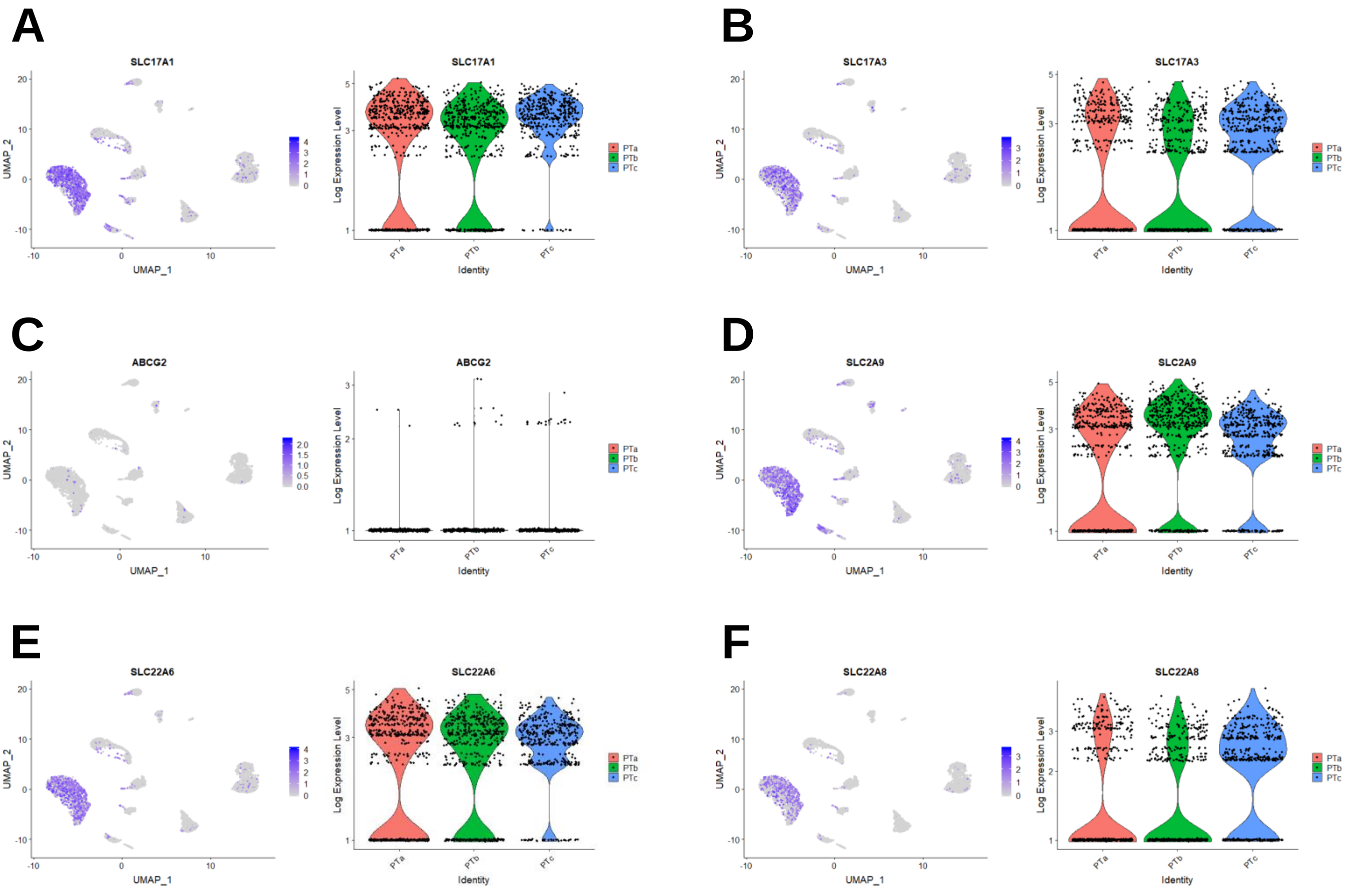

Matsubayashi, Sakaguchi et al. Supplemental Figure 5

A

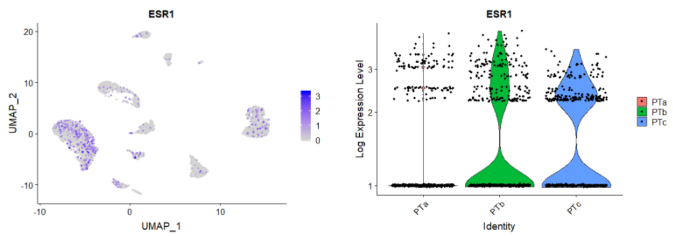

B

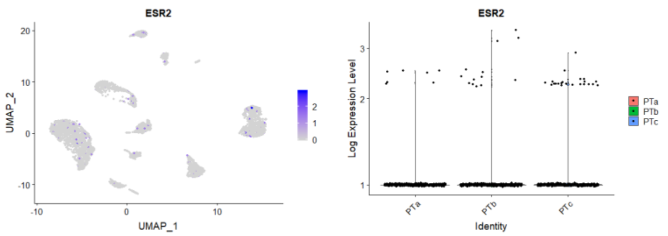

C

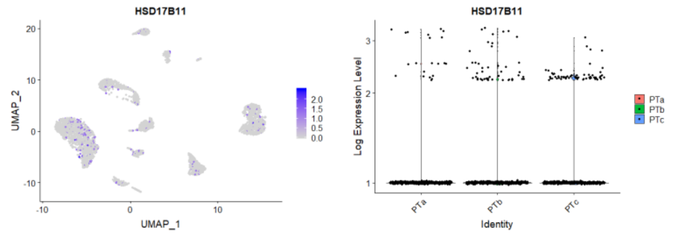

D

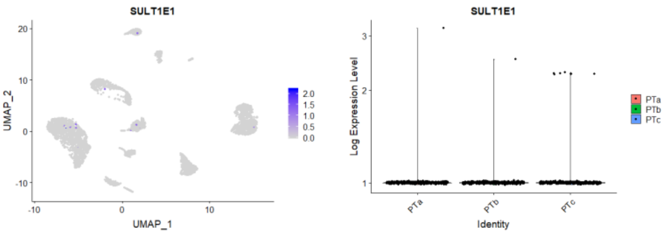

E

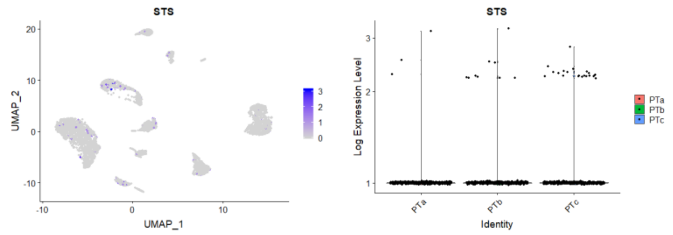

Matsubayashi, Sakaguchi et al. Supplemental Figure 6

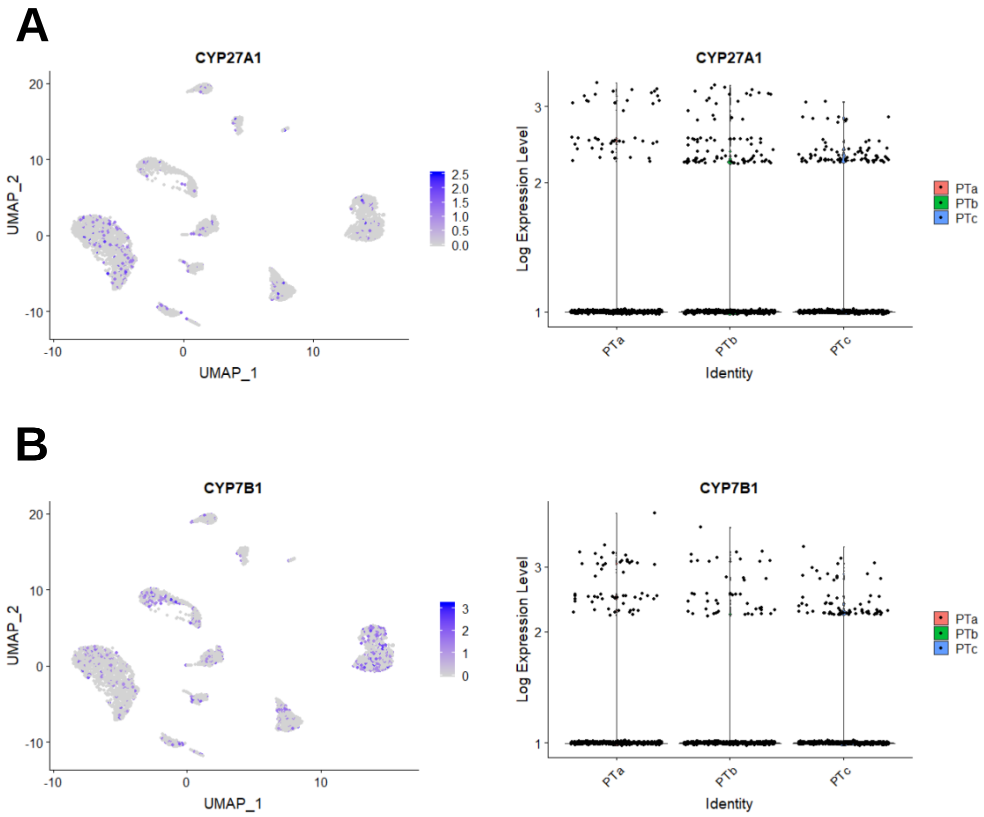

Matsubayashi, Sakaguchi et al. Supplemental Figure 7

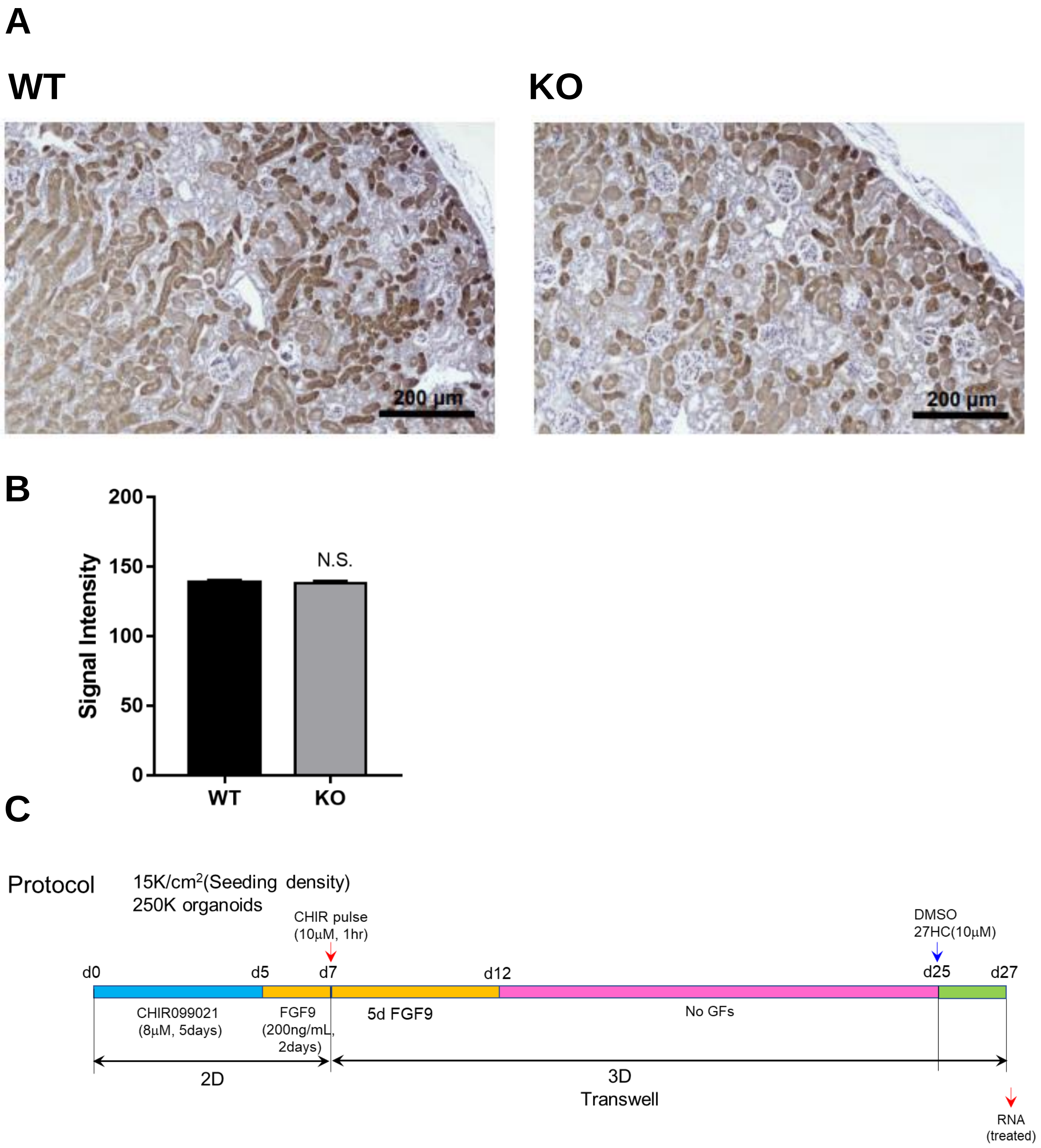

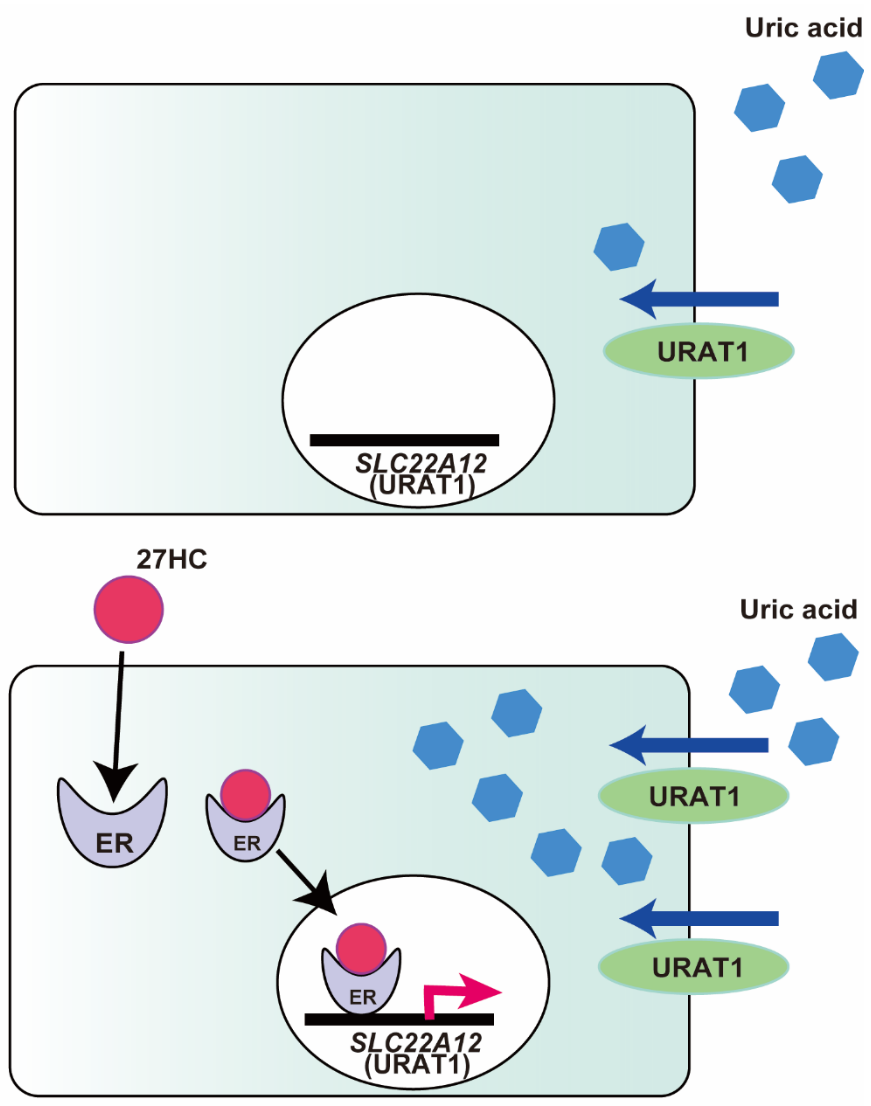
Uric acid
URAT1
SLC22A12
(URAT1)
27HC
Uric acid
ER
ER
URAT1
ER
SLC22A12
(URAT1)
URAT1
